# Proton-pump inhibitors increase *C. difficile* infection risk by altering pH rather than by affecting the gut microbiome based on a bioreactor model

**DOI:** 10.1101/2024.07.10.602898

**Authors:** Julia Schumacher, Patrick Müller, Johannes Sulzer, Franziska Faber, Bastian Molitor, Lisa Maier

**Affiliations:** Cluster of Excellence EXC 2124 Controlling Microbes to Fight Infections, University of Tübingen, Tübingen, Germany; Environmental Biotechnology Group, Department of Geosciences, University of Tübingen, Tübingen, Germany; Interfaculty Institute for Microbiology and Infection Medicine Tübingen, University of Tübingen, Tübingen, Germany; M3-Research Center for Malignome, Metabolome and Microbiome, University of Tübingen, Tübingen, Germany; Helmholtz Centre for Infection Research, Helmholtz Institute for RNA-based Infection Research (HIRI) Würzburg, Germany; Institute for Molecular Infection Biology, Julius-Maximilians-University of Würzburg, Würzburg Germany

**Keywords:** proton-pump inhibitor, gut microbiota, *Clostridioides difficile* infection, bioreactor, colonization resistance

## Abstract

*Clostridioides difficile* infections often occur after antibiotic use, but they have also been linked to proton-pump inhibitor (PPI) therapy. The underlying mechanism—whether infection risk is due to a direct effect of PPIs on the gut microbiome or changes in gastrointestinal pH—has remained unclear.

To disentangle both possibilities, we studied the impact of the proton-pump inhibitor omeprazole and pH changes on key members of the human gut microbiome and stool-derived microbial communities from different donors *in vitro*. We then developed a custom multiple-bioreactor system to grow a model human microbiome community in chemostat mode and tested the effects of omeprazole exposure, pH changes, and their combination on *C. difficile* growth within this community.

Our findings show that changes in pH significantly affect the gut microbial community’s biomass and the abundances of different strains, leading to increased *C. difficile* growth within the community. However, omeprazole treatment alone did not result in such effects. These findings imply that the higher risk of *C. difficile* infection following proton-pump inhibitor therapy is probably because of alterations in gastrointestinal pH rather than a direct interaction between the drug and the microbiome. This understanding paves the way for reducing infection risks in proton-pump inhibitor therapy.

## Introduction

*Clostridioides difficile* has become the most common cause of antibiotic-associated diarrhea, ranging from mild to life-threatening colitis ^1^. Antibiotics favor *C. difficile* infections (CDIs) by disrupting the gut microbiome’s protective barrier, creating an environment that promotes spore germination and *C. difficile* growth. While nearly all classes of antibiotics can increase the risk of CDI, the highest risk is associated with broad-spectrum antibiotics, including clindamycin, fluoroquinolones, and cephalosporins ^2–5^. However, *C. difficile* infections can occur without prior antibiotic use ^6–8^. Other factors, such as the use of proton-pump inhibitors (PPIs), have been shown to increase the risk of CDI in several clinical studies ^9–12^. PPIs are indicated for the treatment of conditions like gastroesophageal reflux disease and ulcers ^13^. As such, they are typically used long-term^14^ and are among the most frequently prescribed drugs worldwide, with omeprazole being the most common ^15^. PPIs inhibit the proton/potassium (H^+^/K^+^)-ATPase enzyme in gastric parietal cells, thereby causing an increase in gastric pH. The reasons why PPI consumption is associated with an increased risk of CDI remain unclear.

PPI consumption, similar to antibiotic use, has been linked to alterations in the gut microbiome composition in various studies ^16–18^. These changes include an increase in *Enterococcaceae*, *Lactobacillaceae*, *Micrococcaceae*, *Pasteurellaceae*, *Staphylococcaeceae*, and *Streptococcaceae*, along with a decrease in *Ruminococcaceae* ^10,16,17^. These findings imply that PPIs, like antibiotics, disturb the protective barrier of the gut microbiome, fostering an environment conducive to CDI. While the direct inhibitory effects of broad-spectrum antibiotics on the gut microbiome have been known for decades ^19^, how PPIs cause changes in the gut microbiome composition remains largely unexplored.

There are two plausible, not mutually exclusive, explanations for how PPIs affect the microbiome: (1) through direct interaction with gut microbes and (2) by their effect on stomach pH. Recent reports indicate that non-antibiotic drugs can directly inhibit members of the gut microbiome, with an estimated 24% of human-targeted drugs inhibiting the growth of key microbiome members ^20^. By directly targeting gut microbes, PPIs could reduce diversity and shift species abundance, explaining the compositional changes observed in microbiome studies.

Alternatively, the effect of PPIs on the microbiome may be a secondary consequence of changes in gastrointestinal pH. Treatment with PPIs, such as omeprazole, raises the gastric pH above 6 and increases the pH of the proximal duodenum ^21,22^. However, the pH-increasing effect diminishes in the distal duodenum, and the pH normalizes when reaching the proximal jejunum ^21,23^. Although the pH change in the stomach is not thought to affect later parts of the intestinal tract, PPI treatment could still lead to a pH change in the colon. Colonocytes express a homolog of the H^+^/K^+^-ATPase found in gastric parietal cells, and omeprazole has been proposed to inhibit also this enzyme. This could increase pH levels within the colon and stool of individuals using PPIs ^21,24–27^. Notably, CDI has been linked to more alkaline stool ^28^. It is tempting to speculate that the more alkaline colon environment created by PPI treatment may promote *C. difficile* growth, thereby increasing the risk of infection.

In this study, we sought to understand how omeprazole influences the microbiome composition to promote *C. difficile* growth. We focused on determining whether these effects are solely mediated by the drug itself or whether changes in pH play a role. In humans and animal models these effects are interconnected and, therefore, difficult to separate. Thus, we employed various *in vitro* systems, ranging from batch cultivation to bioreactor systems, to precisely quantify the consequences of pH changes and physiological omeprazole concentrations on defined and human-stool-derived gut microbial communities. Subsequently, we challenged pH- and drug-perturbed communities with *C. difficile* and monitored its growth within these communities ^29^. Our findings provide strong evidence that the increase in *C. difficile* growth associated with omeprazole is primarily a result of pH changes rather than direct interference of the drug with gut microbes.

## Results

### In monoculture, key members of the human gut microbiome respond to pH change but not to omeprazole

To distinguish between the direct impact of PPIs on the human gut microbiome and the effect of altered gastrointestinal pH, we investigated the PPI omeprazole and the pH sensitivity of key microbiome members. Recognizing that gut microbes are most sensitive to perturbation when grown in monoculture ^30^, which is due to the lack of cross-protection found in communities, we first examined the effects of pH and the drug in monocultures. We selected 21 prevalent and abundant members of the human gut microbiome, which can be studied both in monoculture and as part of a community, referred to as Com21 (Suppl. Table 1). These 21 bacterial species represent 7 bacterial phyla, 11 families, and 18 genera, covering 68.6% of the pathways detected in the human microbiome ^29^.

To assess omeprazole sensitivity, we revisited previous data from our lab ^29^, where bacterial growth in mGAM was measured over 20 hours in the presence of varying omeprazole concentrations. From the same dataset, we also examined the sensitivity of our strains to clindamycin, an antibiotic that is associated with a high risk for CDI ^2–5^. We quantified drug sensitivity by calculating the relative growth in the presence of the drug compared to an untreated control based on the maximum optical density (OD) in the stationary phase. As expected, clindamycin completely inhibited 15 of the 19 tested strains at the lowest concentration of 1.25 µM (Figure 1A).

**Figure 1.**
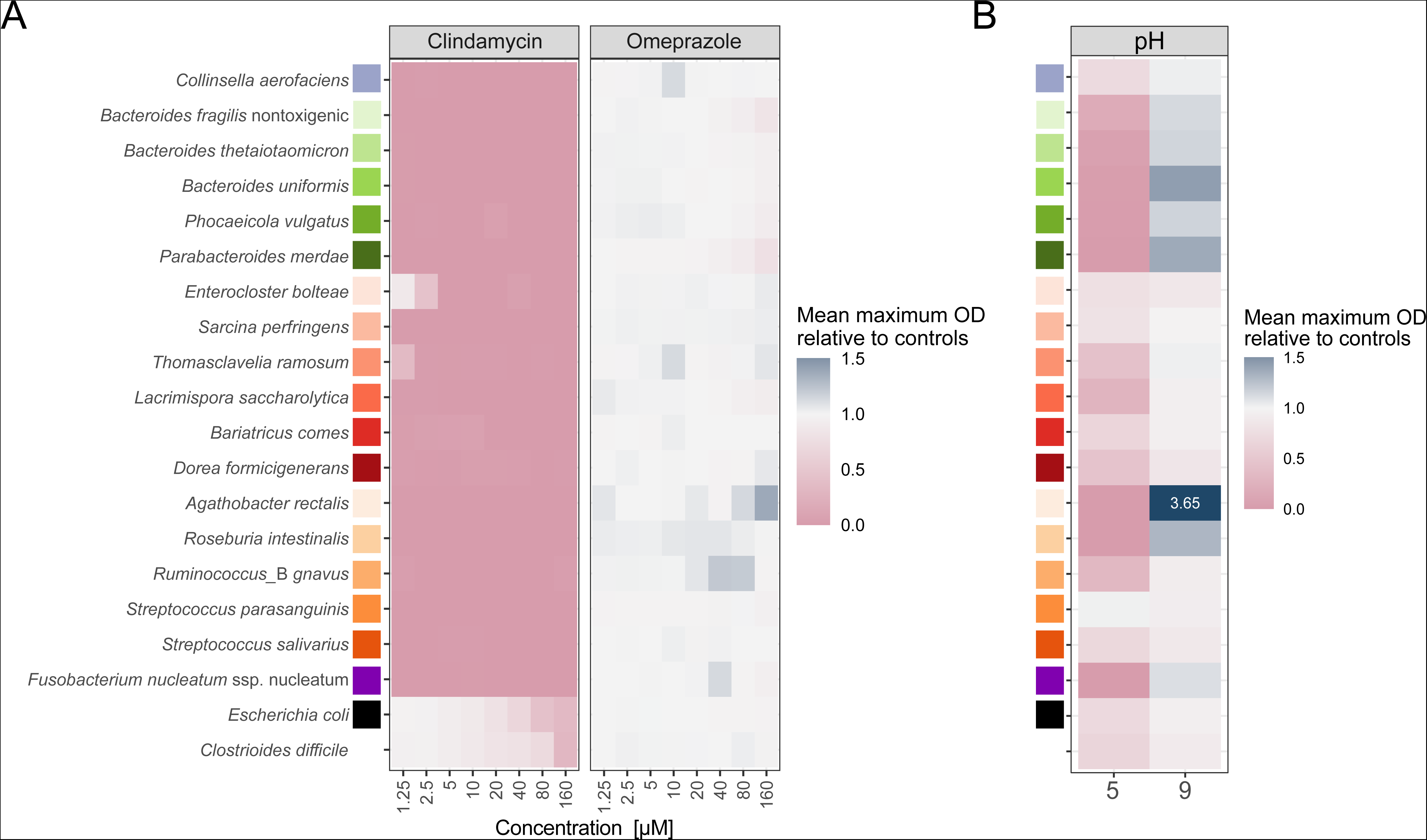
Individual sensitivity of 19 Com21 members to omeprazole and pH. **A)** Growth of Com21 members in the presence of different concentrations of clindamycin and omeprazole in monoculture. Heatmap depicts the mean maximum optical density (OD) of cultures in the stationary phase compared to untreated controls (N = 3). **B)** Growth of Com21 members at different pH in monoculture. Heatmap depicts the mean maximum OD of cultures in the stationary phase compared to OD at pH 7.4. Values outside the legend range are written within the heatmap tile (N = 3).

*C. difficile* showed the highest resistance to clindamycin, maintaining growth with a relative mean OD of 0.71 at 80 µM and 0.31 at 160 µM compared to untreated controls. *Escherichia coli* also tolerated higher clindamycin concentrations, with relative mean ODs of 0.67 at 40 µM and 0.43 at 80 µM. Additionally, *Enterocloster bolteae* (relative OD of 0.87 at 1.25 µM and 0.44 at 2.5 µM) and *Thomasclavelia ramosa* (relative OD of 0.35 at 1.25 µM) were able to grow at the lowest clindamycin concentrations. In contrast, omeprazole did not significantly affect the growth of any of the Com21 members or *C. difficile*; only at 160 µM did some *Bacteroidales* show a slight reduction in OD (relative mean OD 0.78 - 0.9) (Figure 1A). These results indicate that the PPI omeprazole does not directly inhibit commensal bacterial growth. Therefore, unlike clindamycin, the increased risk for CDI associated with omeprazole is likely not due to growth inhibition of gut bacteria.

Next, we investigated the pH sensitivity of the strains. The normal pH in the gastrointestinal tract varies depending on the intestinal site, diet, gender, and health status but generally falls between 5.5 and 7.5 ^21,31–34^. However, it can reach pH 8 to 9 in extreme cases ^33^. We quantified pH sensitivity at pH 5 and pH 9 by measuring the maximum OD in the stationary phase and normalizing it to growth at pH 7.4. Overall, pH 5 had a more severe impact on growth, with several species being unable to grow at this pH (e.g., all *Bacteroidales*; Figure 1B). Only *Streptococcus parasanguinis* was unaffected at pH 5. Growth at pH 9 was less impaired, with all strains being able to grow. Some species, such as the *Bacteroidales*, *Agathobacter rectalis*, and *Roseburia intestinalis*, even grew better at pH 9 compared to pH 7.4 (Figure 1B).

Our data shows that pH sensitivity varies across species, with lower pH (pH 5) more severely inhibiting bacterial growth. This underscores the importance of low pH in the upper small intestine and stomach as a barrier to incoming bacteria—a barrier that is compromised during long-term PPI treatment, potentially leading to small intestinal bacterial overgrowth ^13^. At pH 9, we observed some growth improvement but generally more growth impairment, indicating that pH changes can affect community structure, abundance, and functions. Thus, our results in monocultures suggest that pH has a more severe impact on individual community members than omeprazole does.

### Batch-cultivated human stool-derived communities are insensitive to omeprazole and subsequent *C. difficile* challenge

Monocultures of selected gut bacterial species fail to fully capture the species diversity, interspecies variation, and individual compositional differences that characterize gut microbiomes. Therefore, we examined microbial communities derived from human fecal samples of healthy donors to assess their sensitivity to omeprazole. Subsequently, we exposed omeprazole-treated communities to *C. difficile* and quantified its growth in these communities.

We first revisited previous data on the omeprazole sensitivity of human fecal samples ^29^. Consistent with single-species sensitivities, the growth of all stool-derived communities was unaffected by omeprazole at all tested concentrations (Figure 2A). In contrast, clindamycin sensitivity varied between donors. For example, the community from human fecal sample 5 still grew to a relative mean OD of 0.54 at the highest clindamycin concentration of 160 µM. In comparison, the community from human fecal sample 7 was already reduced to a relative mean OD of 0.46 at the lowest concentration of 1.25 µM (Figure 2A). This highlights the inherent differences in antibiotic sensitivities among human gut microbiomes.

**Figure 2.**
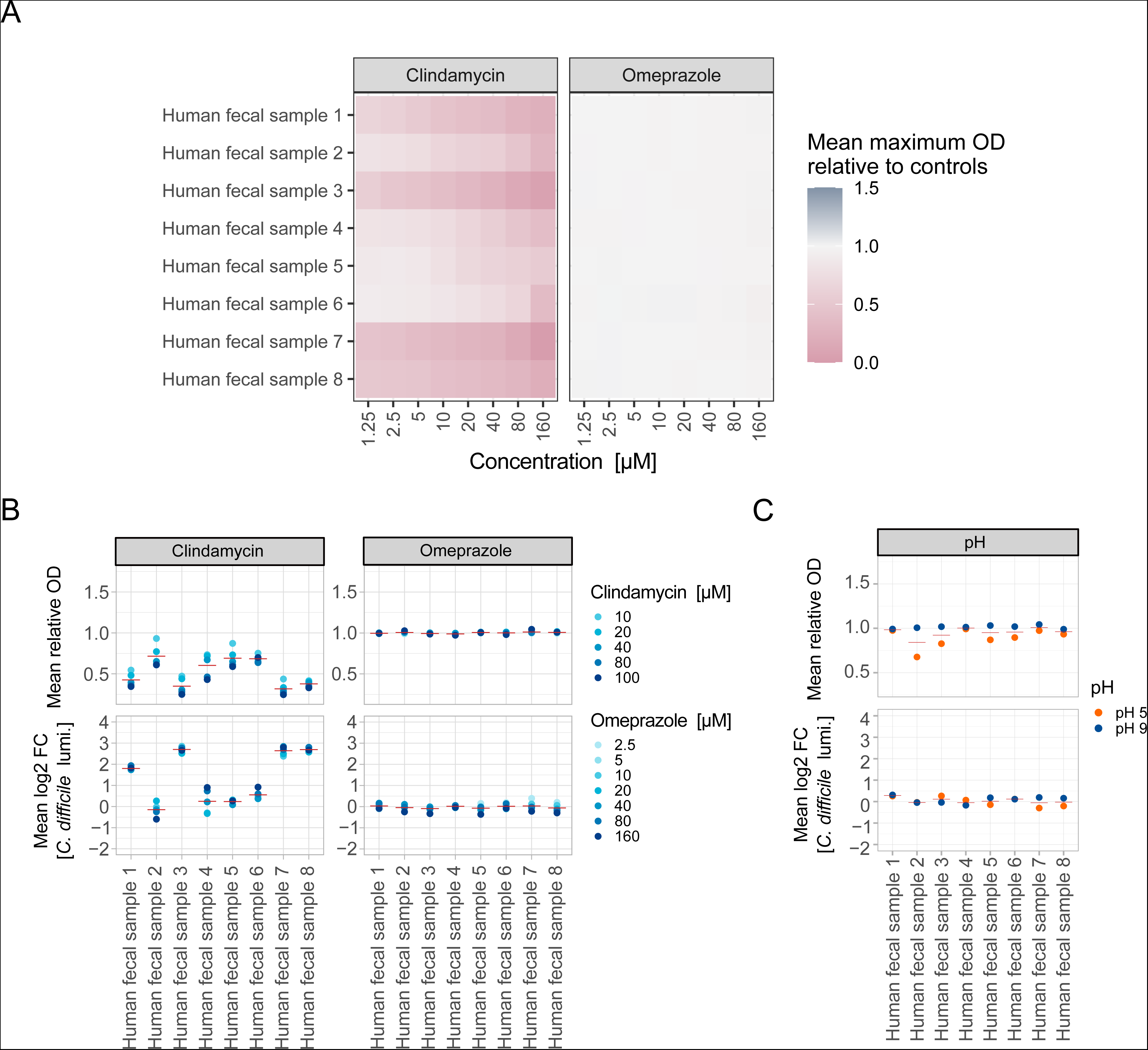
Neither omeprazole treatment nor changes in pH promote the growth of C. difficile within human stool-derived microbial communities. **A)** Growth of communities derived from eight human fecal samples in the presence of different concentrations of clindamycin (left) and omeprazole (right). Heatmap depicts the mean maximum optical density (OD) of cultures in the stationary phase compared to untreated control growth (N = 3). **B) Top:** Mean OD of the eight communities relative to untreated controls after treatment with different concentrations of clindamycin (left) or omeprazole (right) for 24 h. Red horizontal line depicts the mean per fecal sample across all concentrations (N = 3) **Bottom:** Mean log2 fold change (FC) of *C. difficile* growth as determined by *C. difficile* luminescence after 5 h in clindamycin (left) or omeprazole (right) treated communities relative to untreated controls. Red horizontal line depicts the mean per fecal sample across all concentrations (N = 3). **C) Top:** Mean OD of eight human stool-derived communities relative to controls at pH 7.4 after growth at different pH for 24 h. Red horizontal line depicts the mean per community (N = 3) **Bottom:** Mean FC of *C. difficile* growth as determined by *C. difficile* luminescence after 5 h in pH-exposed communities relative to controls at pH 7.4. Red horizontal line depicts the mean per fecal sample (N = 3).

Furthermore, we investigated whether omeprazole exposure affects *C. difficile* growth within stool-derived communities. We exposed the communities to various concentrations of omeprazole (2.5 µM to 160 µM) for 24 hours before challenging them with *C. difficile* carrying a constitutive plasmid-based luminescence reporter (Extended Data Figure 1A). The untreated stool-derived communities were able to significantly reduce *C. difficile* growth after 5 hours compared to *C. difficile* grown in monoculture, as measured by luminescence (mean relative *C. difficile* growth in communities: 1.79% ± 0.1 standard error of the mean (SEM)) (Extended Data Figure 1B). Consistent with the omeprazole sensitivity data, the biomass/OD of the community after 24 hours of omeprazole exposure did not change relative to unperturbed controls. Additionally, omeprazole did not impact *C. difficile* growth (Figure 2B, bottom right).

As a positive control, we conducted the same challenge with clindamycin-treated stool-derived communities (10 µM to 100 µM). Clindamycin reduced community biomass to varying degrees among donors in a concentration-dependent manner. Consistent with clinical observations, clindamycin also increased *C. difficile* growth after pathogen challenge in at least four of the eight samples, up to 9-fold (Figure 2B, bottom left). Overall, communities that were more sensitive to clindamycin exhibited higher levels of *C. difficile*, indicating that reduced biomass correlates with increased *C. difficile* growth (Pearson’s ρ = −0.8612, p-value = 9.986e-13).

We also investigated the effect of pH changes on stool-derived fecal communities. The communities were grown at pH 5, pH 9, or physiological pH 7.4 for 24 hours before being challenged with *C. difficile*. Growth at pH 5 reduced the biomass of some fecal communities to a minimum relative OD of 0.68 (Figure 2C, top). However, neither pH 5 nor pH 9 affected subsequent *C. difficile* growth within the communities (Figure 2C, bottom).

These results indicate that neither the PPI omeprazole nor pH changes directly affected *C. difficile* growth in stool-derived communities from different donors. However, it is important to note that both omeprazole and pH exposure were limited to 24 hours, with pH adjustments made only at the beginning of the experiment. Since PPIs are typically used long-term, more prolonged exposure to the drug and sustained pH changes would need to be investigated.

### Multiple-bioreactor system enables precise studies of microbiome perturbations

To overcome the limitations of working with stool-derived communities in batch, we turned to chemostats and our gut model community Com21. Chemostats allow precise, continuous adjustment and monitoring of environmental conditions, such as pH, over long periods, making them the ideal system to address our question. We chose a previously described system ^35^ based on six bioreactor bottles (Figure 3A and 3B), which can be operated simultaneously, individually at different conditions, or as replicates. Our multiple-bioreactor system (MBS) can be operated under an aerobic or anaerobic atmosphere and can be used for batch cultivation or in chemostat mode, where fresh medium is continuously supplied and spent medium is removed at the same rate.

**Figure 3.**
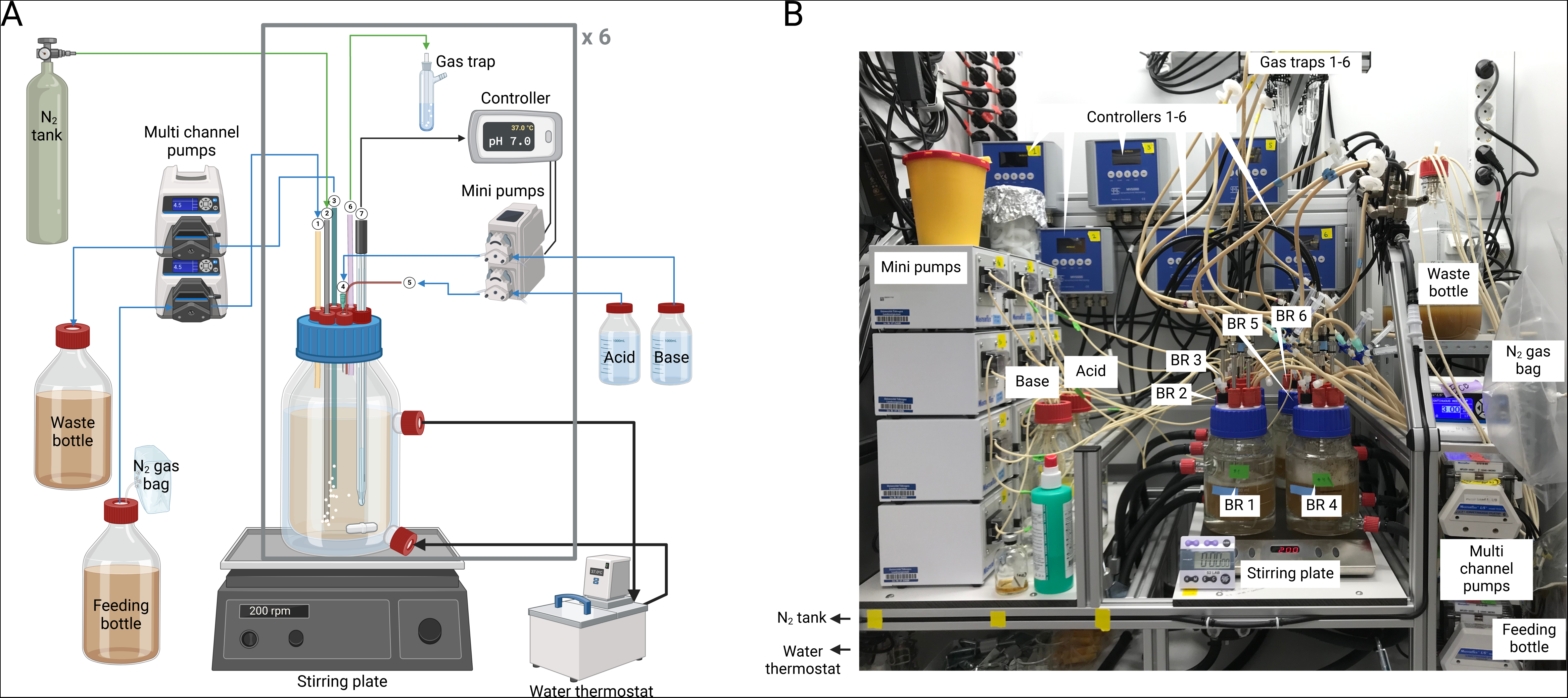
Overview of the multiple-bioreactor system. **A)** Schematic overview of a single bioreactor bottle. Each double-walled bioreactor bottle has a volume of 500 mL. 1: Fresh medium is introduced at a desired flow rate through the feeding port from a feeding bottle; 2: The desired gas mix is introduced into the bottle through a stainless steel tube with an attached sparger; 3: Spent medium is removed through a stainless steel tube. This port is also used as a sampling port where samples can be taken with a syringe; 4: A syringe punched through a rubber stopper is used as base port; 5: The acid port is a button once cannula punched through the same rubber stopper as the base port; 6: The gas-out port is connected to a foam trap; 7: An autoclavable pH/pt1000-electrode measuring the pH and temperature is connected to a multi-parameter controller. Each controller is connected to two mini-pumps that are triggered to pump acid or base when the pH falls out of range. All bioreactor bottles are placed on a multi-stirrer plate and stirred with a magnetic stirrer. Temperature is maintained by a water jacket connected to a water thermostat. As indicated by the brackets and the x6 we have 6 bioreactors which can be used simultaneously. Created with BioRender.com. **B)** Picture of the complete setup. BR: bioreactor.

To demonstrate the functionality and reproducibility of our MBS with a gut microbial community, we conducted a pilot experiment in three bioreactors using a reduced version of Com21 ^29^, referred to here as Com18. Com18 lacks *E. coli*, *Veillonella parvula*, and *Eggerthella lenta* due to concerns about *E. coli* dominance and initial difficulties in growing *Veillonella* and *Eggerthella* species. The MBS was operated anaerobically for 188.75 hours, starting with 24 hours of batch mode followed by continuous operation in chemostat mode. The OD increased during continuous operation until the 50-hour mark, after which it stabilized (Extended Data Figure 2A). We noted fluctuations in the bioreactor volumes among the individual replicates, which might explain the variations in OD. The pH remained stable at 7 (± 0.259) throughout the experiment. The community composition was determined by 16S rRNA gene sequencing (Extended Data Figure 2B).

During the first few hours of batch mode, the community was dominated by S*arcina perfringens* and *Streptococcus salivarius*. By the end of the batch phase at 22 hours, the abundance of *Fusobacterium nucleatum*, *Bacteroides thetaiotaomicron*, and *Phocaeicola vulgatus* increased. During continuous operation, *Bacteroides uniformis* and *F. nucleatum* further increased in abundance, while *S. perfringens*, *P. vulgatus*, *R. intestinalis*, *S. salivarius*, and *S. parasanguinis* decreased, despite the overall biomass (OD) remaining constant. *R. intestinalis* and *S. salivarius* were lost in all replicates at 116.75 hours and 22 hours, respectively. *A. rectalis* and *Ruminococcus gnavus* were only detected in one replicate at the final time point and were absent in the other two replicates. No contamination with non-Com18 species was observed with sequencing.

The community composition and biomass of all three replicates were similar. This pilot experiment demonstrated the suitability of our MBS and evaluation methods for studying microbial communities under controlled conditions.

### In the MBS, pH changes promote *C. difficile* growth in Com21, whereas omeprazole treatment does not

Next, we used the MBS with Com21 to investigate the effects of pH changes or exposure to omeprazole on the subsequent growth of *C. difficile* (Figure 4). We allowed Com21 to stabilize for six days, which corresponds to six hydraulic retention times (HRTs), before adjusting the pH to 5 (bioreactors 1 and 4), 9 (bioreactors 3 and 6), or 7 (bioreactors 2 and 5). Following an additional six days at the respective pH levels, five out of the six bioreactors (excluding control bioreactor 5) were treated with 80 µM omeprazole for three consecutive days (every 24 hours), based on its estimated concentration in the human intestine ^20^. Afterwards, all bioreactors were set to recover at pH 7 for six days. Samples for the pathogen challenge assay and 16S rRNA gene sequencing were collected after stabilization, pH changes, each day of omeprazole treatment, and after recovery, totaling six samples per bioreactor (Figure 4A). At each sampling point, samples were challenged with *C. difficile* using the same assays described for the stool-derived communities.

**Figure 4.**
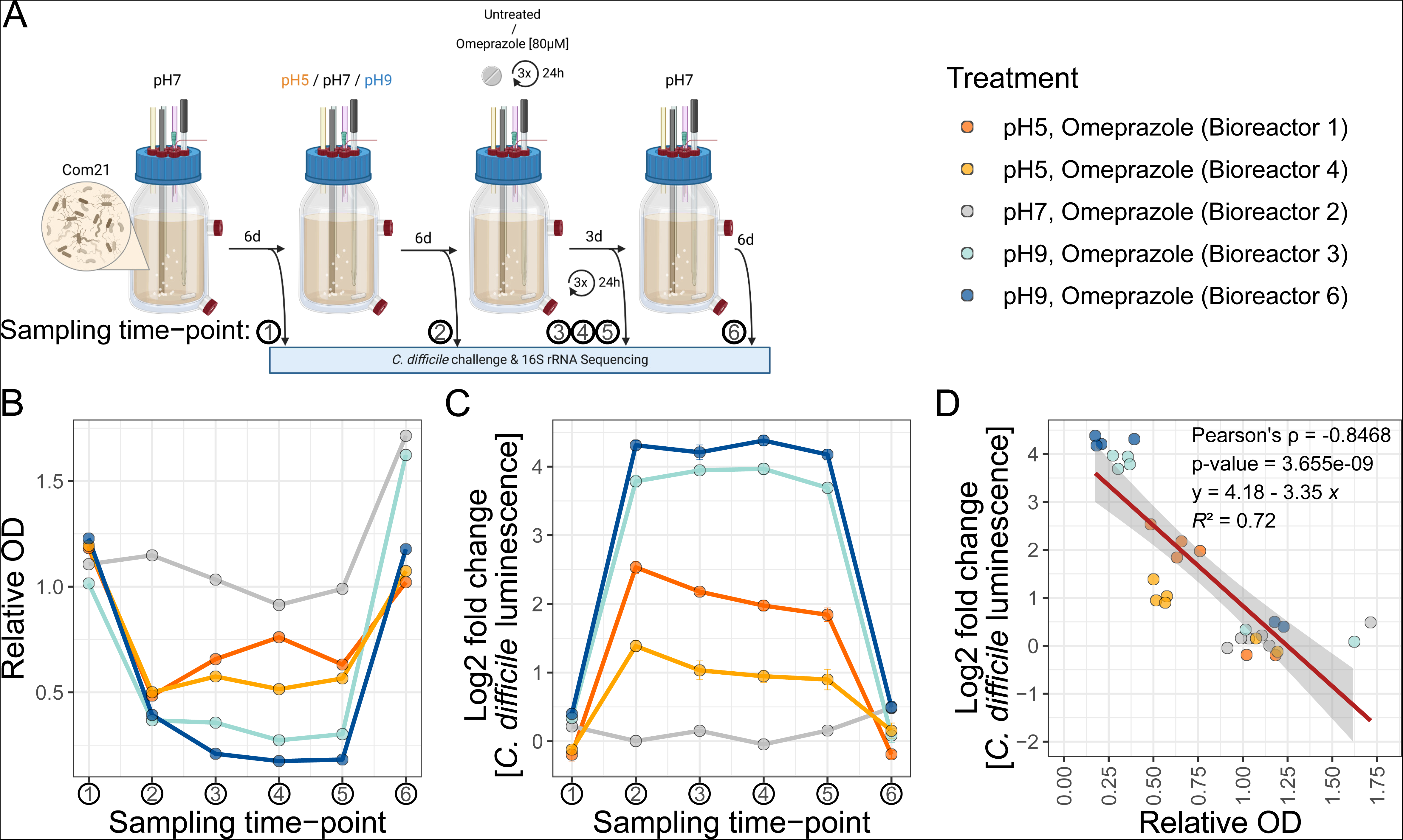
Change in pH decreases community biomass and increases growth of *C. difficile*. **A)** Schematic overview of the bioreactor workflow. Com21 was grown for six HRTs in chemostat mode with mGAM at pH 7. After this period, the pH was either changed to pH 5 or pH 9 for six HRTs or left unchanged. Subsequently, omeprazole was added daily at 80 µM to five of the six bioreactors for three consecutive HRTs, after which all bioreactors were returned to pH 7 for other six HRTs. Sampling points are indicated with arrows. Created with BioRender.com. **B)** Relative OD of the bioreactors over time at every sampling time point compared to the median OD of the untreated control (bioreactor 5). Bioreactors 1 and 4 were switched to pH 5, bioreactors 3 and 6 to pH 9, and bioreactor 2 remained at pH 7. All bioreactors, except the control bioreactor 5 used for normalization, underwent omeprazole treatment at 80 µM. **C)** Log2 fold change in *C. difficile* growth in the bioreactor communities at every time point. *C. difficile* growth was quantified by luminescence measurement after 5 h and normalized to the median luminescence of *C. difficile* in the untreated control (bioreactor 5) at the same time point. The mean with standard deviation of 11 technical replicates is shown. **D)** Correlation of relative community OD to log2 fold change in *C. difficile* growth. Values from plots B and C are shown with colors indicating the corresponding bioreactor. The red line represents the linear trendline, with its function, *R*² value, Pearson correlation, and p-value provided in the plot.

The untreated Com21 community from bioreactor 5 strongly inhibited *C. difficile* growth to levels observed for stool-derived communities (mean relative *C. difficile* growth in Com21: 1.58% ± 0.19 SEM) (Extended Data Figure 1B). Initially, after six days at pH 7, all bioreactor communities exhibited comparable biomass and similar protection against *C. difficile* challenge, which remained consistent throughout the 21-day experiment in the control bioreactor (Figure 4). When bioreactor 2’s community, maintained at pH 7, was exposed to 80 µM omeprazole, there was no observable change in OD or *C. difficile* growth, indicating that omeprazole did not directly impact Com21 in our setup (Figure 4B and C). This is in line with our observation that omeprazole did not strongly affect the growth of Com21 members in monocultures (Figure 1A).

However, altering the pH of the bioreactors to either pH 5 (bioreactors 1 and 4) or pH 9 (bioreactors 3 and 6) resulted in reduced biomass and increased *C. difficile* growth in those communities compared to the control (up to a 22-fold increase in *C. difficile*, Figure 4B and C). Notably, omeprazole treatment of communities in altered pH did not further impact their biomass or *C. difficile* growth. Overall, we observed a negative correlation (Pearson’s ρ = −0.8468, p-value = 3.655e-09) between the biomass of the community and the growth of *C. difficile* in this community (Figure 4D), similar to what we observed for clindamycin in stool-derived communities in Figure 2B.

The changes in pH were accompanied by strong shifts in microbial community composition, while omeprazole treatment alone did not induce any changes (Figure 5A). Both pH 5 and pH 9 resulted in decreased levels of *F. nucleatum* and *B. uniformis*, the two most dominant species in the bioreactor communities (Figure 5B). Communities at pH 5 showed increased levels of *A. rectalis* and either *Collinsella aerofaciens* (bioreactor 1) or *B. thetaiotaomicron* (bioreactor 4), whereas communities at pH 9 exhibited increased levels of either *B. thetaiotaomicron* and *V. parvula* (bioreactor 3) or *E. lenta* and *T. ramosa* (bioreactor 6). *A. rectalis* was initially absent from all bioreactors but was detected in pH 5-treated bioreactors, where it persisted in low amounts after recovery. Similarly, *S. perfringens* was initially present in low amounts in only one bioreactor but appeared in both pH 5 bioreactors and one pH 9 bioreactor (Figure 5B).

**Figure 5.**
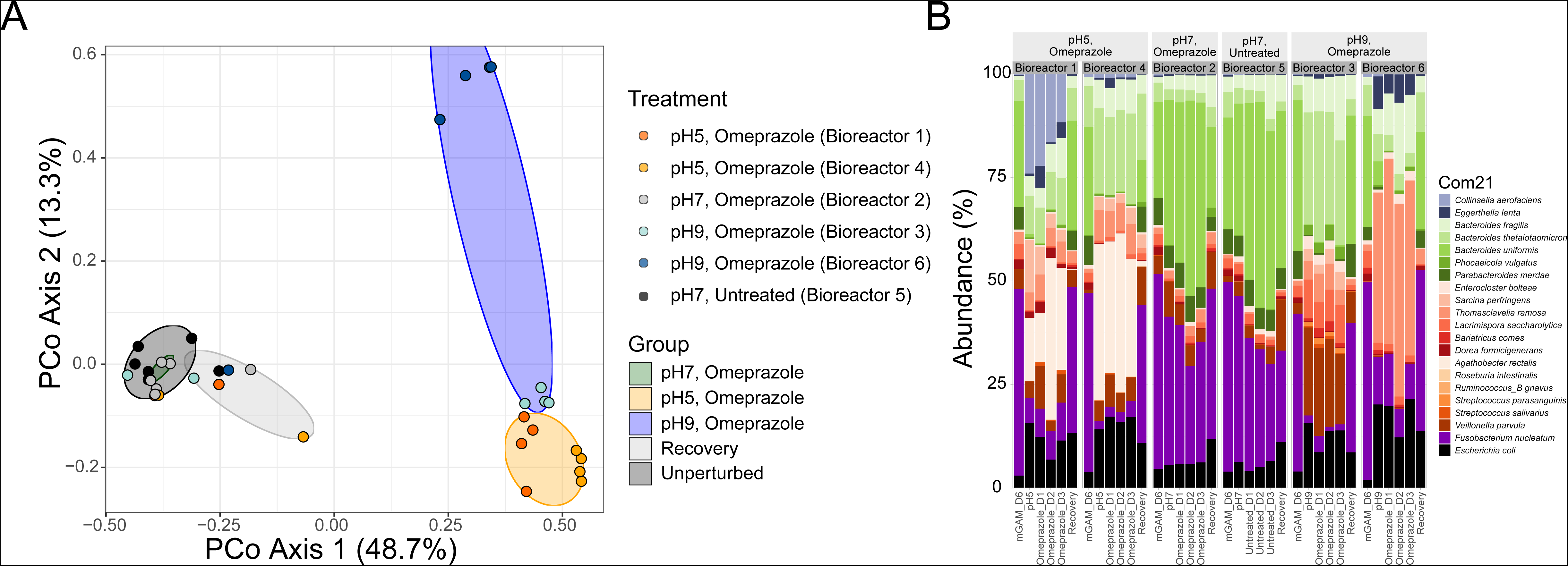
Change in pH causes significant changes in community composition. **A**) Principal Coordinate Analysis of Bray-Curtis dissimilarity. Data points are color-coded by bioreactor and grouped by treatment (colored ellipses). The untreated control (bioreactor 5), initial compositions of all bioreactors, and pH 7 treatment of bioreactor 2 are grouped as ‘Unperturbed’. The three Omeprazole treatment sampling points of bioreactor 2 are grouped as ‘pH7, Omeprazole’. Sampling points at pH 5 (with and without omeprazole) for bioreactors 1 and 4 are grouped as ‘pH 5 Omeprazole’. Sampling points at pH 9 (with and without omeprazole) for bioreactors 3 and 6 are grouped as ‘pH 9 Omeprazole’. All recovery sampling points (except bioreactor 5) are grouped under ‘Recovery’. **B**) Relative abundance of each member of Com21 at the indicated sampling time points. Panels are grouped by bioreactor: Bioreactors 1 and 4 were changed to pH 5 with omeprazole, bioreactors 3 and 6 were changed to pH 9 with Omeprazole, bioreactor 2 was treated with Omeprazole, and bioreactor 5 was left untreated.

These changes in composition were only partially explained by the individual pH sensitivities of the strains (Figure 1B). For instance, *Bacteroidales* were found to be acid sensitive, and indeed, their relative abundance decreased in the bioreactors at pH 5. Conversely, *S. perfringens* showed relatively greater resistance to acidity compared to other members, resulting in its increased relative abundance in the pH 5 bioreactors. However, strains sensitive to acidity, such as *A. rectalis*, increased in the pH 5 bioreactors but not in the pH 9 bioreactors, despite demonstrating substantially better growth at pH 9 in monoculture. Additionally, *S. parasanguinis,* which exhibited acid resistance, was absent from all bioreactors initially and did not increase at pH 5.

Remarkably, all bioreactor communities reverted to their original biomass, resistance against *C. difficile* growth, and community composition after recovery at pH 7 for six HRTs (Figure 4 and 5). In summary, these findings indicate that a shift in pH modifies the composition of the human gut microbial community, resulting in decreased biomass and reduced resistance to *C. difficile* growth. Importantly, omeprazole does not induce such changes on its own, suggesting that the reported association between PPI usage and an increased risk of CDI could be attributed to the prolonged alterations in the pH of the gastrointestinal tract caused by the drug rather than its direct interaction with gut microbes.

## Discussion

CDI often occurs after antibiotic treatment, but the use of PPIs has also been strongly linked to CDI. It was previously unclear whether this link was due to a direct effect of PPIs on *C. difficile* or the microbiome, or if it resulted from altered gastrointestinal pH as a secondary effect of PPIs acting on the host. Disentangling the direct effect of the drug from the secondary pH effect is impossible in *in vivo* models or cohort studies. To address this, we used an MBS to separate the effects of the PPI omeprazole from the effects of altered pH on the gut microbial community. Our results showed that omeprazole does not directly affect the composition of a synthetic community of human gut commensals or its ability to limit the growth of *C. difficile*. In contrast, changes in pH were strongly correlated with altered community compositions, reduced biomass, and, ultimately, increased growth of *C. difficile* in pH-perturbed communities. Thus, our data support the hypothesis that PPIs increase the risk for CDI not by direct drug-microbe interaction but by changing the pH of the gastrointestinal tract ^16^.

In monocultures, we observed that only some *Bacteroidales* showed slight sensitivity to omeprazole. However, this did not result in lower *Bacteroidales* levels in omeprazole-treated MBS communities. This contrasts with clinical studies that reported a decrease in *Bacteroidetes* after omeprazole treatment linked to CDI ^9,21^. Therefore, our results suggest that the reduction in *Bacteroidetes* seen in patients may be due to factors other than the drug’s direct inhibition.

Environmental factors, such as pH, strongly impact community composition due to the varying pH sensitivities of different species. In this study, we deliberately used pH levels at the extremes of what might be expected in a clinical context. This was done to mimic the potential increase in gastrointestinal pH due to PPI administration and to fully assess the impact of these pH changes on *C. difficile* growth in communities. These pH shifts can indicate which species are more resilient to such changes. For example, *C. aerofaciens*, known for its acid tolerance^36^, thrived in our pH 5 communities. Conversely, and consistent with our findings on pH sensitivity in monocultures, *Bacteroides* species showed acid sensitivity, decreasing in numbers in our bioreactor experiments ^9,36,37^. This aligns with clinical reports of decreased *Bacteroidetes* following PPI treatment in patients ^9,21^, suggesting that the association between PPIs and CDI is due to pH changes rather than a direct effect of omeprazole.

Furthermore, species such as *A. rectalis*, which we found to be acid-sensitive in monoculture, were able to thrive in our pH 5 bioreactors. These species have also been previously shown to increase in bacterial communities at pH 5.5 ^34^. The seemingly contradictory discrepancy in pH sensitivity of monocultures versus bioreactor communities highlight that the presence and abundance of certain species in a community cannot be inferred solely from their individual sensitivity *in vitro*. These findings underscore that bacterial community properties are emergent and cannot be fully explained by the sum of individual characteristics, such as pH sensitivity.

The gut microbiota protects against *C. difficile* through various mechanisms. These include producing inhibitory metabolites, such as secondary bile acids, short-chain fatty acids (SCFAs), and antimicrobials, as well as competing for nutrients, particularly proline and other amino acids essential for Stickland fermentation ^13,38^. In our bioreactors, altered pH resulted in reduced biomass and changes in species abundances, such as lower levels of *Bacteroidetes*, potentially creating niches for *C. difficile*. However, associating specific species or strains with increased or decreased resilience against *C. difficile* is challenging because even in our controlled bioreactor setup, we observed different microbiome shifts when applying the same pH shift. Thus, microbiome disturbances do not always result in the same effects at the single species level but are more apparent on broader community properties, such as impairing *C. difficile* growth. This shows that various deviations from the physiological microbiome composition can lead to increased pathogen susceptibility, underlining the importance of systems-based approaches to understand interactions at the microbial community and microbe-host levels.

With increasing knowledge about microbiomes and their importance for human health, there is a pressing demand for approaches to study them effectively. Although bioreactors constitute simplified models, they allow for the detailed study of specific aspects by breaking down the complex system into individually controllable parameters. Thus, bioreactor studies facilitate a deeper understanding of biological processes. While commercial bioreactor systems are expensive and require expertise from manufacturing companies for setup and operation, cost-effective and off-the-shelf alternatives are available ^35,39,40^. These setups can enhance early drug discovery studies by providing systematic approaches to continuously monitor a drug’s effect on gut microbial communities over an extended period, either preceding or accompanying *in vivo* models or clinical trials.

Of note, our study is limited by the inability to investigate host contributions relevant to CDI risk within our *in vitro* systems. These include aspects of the innate and adaptive immune responses, the host’s metabolism, as well as other host-derived factors that influence the *C. difficile* cycle, such as the enterohepatic circulation of bile acids and their interaction with the microbiome ^41^. Our approach also does not account for the direct effects of pH changes or omeprazole exposure on *C. difficile* virulence. Specifically, PPIs and non-physiological pH levels have been reported to increase *C. difficile* toxin expression ^42^, which is crucial for inducing colitis in patients. Thus, to thoroughly determine whether the PPI-mediated increased risk of CDI is due to potential direct interactions of omeprazole with the gut microbiome or a pH-shift dependent mechanism, further investigations are needed, including an exploration of more subtle pH changes. Ultimately, such research will enhance our understanding of microbiome-mediated side effects of PPIs, opening up broad possibilities for mitigating these effects and improving drug safety.

## Methods

### Bacterial cultivation

The species and strains used in this study can be found in Supplementary Table 1. They were purchased from DSMZ and ATCC or were a gift from the Denamur Laboratory (INSERM). In the present manuscript, we use the taxonomic classification from the genome taxonomy database (GTDB) release R06-RS202.

Bacterial cultivation in monoculture was conducted as described before ^43^. In brief, all species were cultivated in mGAM medium (HyServe GmbH & Co.KG, Germany) at 37°C except for *V. parvula*, which was grown in Todd-Hewitt Broth supplemented with 0.6 weight-% sodium lactate. The plasmid-carrying *C. difficile* strain (LM0061) was cultivated in mGAM with 15 µg/mL thiamphenicol. All media, glass, and plastic ware were pre-reduced for a minimum of 24 h under anaerobic conditions (2 vol-% H_2_, 12 vol-% CO_2_, 86 vol-% N_2_) in an anaerobic chamber (Coy Laboratory Products Inc.). Species were inoculated from frozen glycerol stocks into liquid culture medium and passaged twice (1:100) overnight before being used in subsequent experiments. To ensure no contamination of species occurred, their purity and identities were regularly checked *via* 16S rRNA-gene sequencing and/or MALDI TOF mass spectrometry (MS) ^44^.

Stable communities from human fecal samples^45,46^ were inoculated from frozen glycerol stocks into liquid culture medium (mGAM) and incubated at 37°C overnight before being used in downstream assays.

Bacterial cultivation in bioreactors was conducted with mGAM medium. Fresh mGAM medium for initial inoculation was sterilized directly in each bioreactor bottle to ensure sterility of all tubings and ports. Upon sterilization, the bioreactor bottles were sparged with N_2_ gas for at least 12 h to achieve anaerobic conditions. No growth (change in OD) after overnight incubation of the medium under a 100 vol-% nitrogen atmosphere at pH 7 and 37°C further confirmed the sterility of the system.

Each species was incubated as described above in monocultures to assemble Com18 or Com21 to inoculate the bioreactors. Afterward, the OD_600_ of every species was measured (Thermo Scientific^TM^ BioMate^TM^ 160 UV-Vis Spectrophotometer), and they were first combined at equal OD_600_ to a final OD_600_ of 0.01 so that every species contributed 0.000556 OD_600_ (Com18) or 0.000476 OD_600_ (Com21) to the culture. The bioreactors were operated in batch mode for the first 24 h to allow sufficient microbial growth. After 24 h the system was switched to continuous mode.

### pH sensitivity of individual Com21 members, C. difficile LM0061, and human fecal samples

To assess pH sensitivity of the community members and the plasmid-carrying *C. difficile* strain (LM0061), bacteria were grown in mGAM for two subsequent overnight cultures as described above. Human fecal samples were grown in mGAM for one overnight culture. Sensitivity to pH was investigated for 19 out of the 21 community members. *E. lenta* is a slow grower with poor growth in mGAM monoculture, and *V. parvula* has different media requirements in monoculture. Thus, neither was analyzed in this assay. The pH of mGAM medium was adjusted to pH 5 with hydrochloric acid, to pH 9 with sodium hydroxide, or left unchanged at pH 7.4. The pH-adjusted media were transferred to sterile Nunclon 96-well U-bottom microplates (Thermo Scientific, cat. no. 168136) inside the anaerobic chamber, and prereduced for at least 24 h. In addition to 95µl of medium, each well was inoculated with 5 µL bacterial culture to a final OD_578_ of 0.01. Growth was measured in a plate reader over 20 h as described before ^43^ and quantified by taking the maximum OD_578_ during the stationary phase and normalizing it to maximum OD_578_ at pH 7.4.

### Bioreactor handling and operating conditions

A list of all the equipment for the construction of the bioreactor system is summarized in Supplementary Table 2. The 500-mL bioreactor bottles were operated with a working volume of 250 mL. The bioreactors were continuously sparged with N_2_ at approximately 2.5 mL min^-1^ to minimize the risk of O_2_ intrusion into the system. Agitation by a magnetic stirrer was set to 200 rpm. The cultivation temperature was set to 37°C (± 0.2 °C) and maintained at any time by a water thermostat circulating water through the double-walled bioreactor bottles. The starting pH was 7 (± 0.05), and the hysteresis was set to 0.01. The pH probes in each bioreactor bottle were calibrated before autoclaving. The pH was measured daily with an external pH probe (pH-electrode pHenomenal® LS 221). For pH control, we used 0.5 M acid and base solutions (the response of the pH probe and the pumps were too slow for molarities above 0.5 M). The bioreactors were inoculated with a starting OD_600_ of 0.01, such that each strain equally contributed to the starting OD_600_. Upon inoculation, the bioreactors were operated in batch mode for 24 h before switching to continuous mode. For continuous mode, we used a medium feed rate of 0.1736 mL min^-1^, corresponding to an HRT of 24 h. Samples to measure the OD_600_ were taken at least every second day. Samples with an OD_600_ higher than 0.5 were diluted 1:10 prior to the measurement. The pH was continuously monitored by the internal pH probe connected to a controller, which would trigger acid or base inflow if the pH deviated ±0.05 from 7. The temperature and the working volume were manually monitored regularly. We observed fluctuations in the bioreactor volumes in continuous mode. Those fluctuations arise from either a medium inflow faster than the outflow or *vice versa*. Small variations in the flow rate for each bioreactor occur due to differences between the cassettes on the pump head. Variations in the flow rate are likely to affect the OD by either diluting out the bacteria or providing more nutrients for faster growth, which becomes visible through changes in the OD_600_. For the omeprazole treatment of the Com21, we dissolved 6.908 mg/ml omeprazole (TCI, CAS 73590-58-6) in DMSO, and added 1 ml of the solution to each bioreactor, resulting in an overall concentration of 80 µM, except for the control reactor, which was treated with 1 ml DMSO. For the *C. difficile* invasion assays, we took a 2-mL sample of each bioreactor with a syringe and directly transferred the samples to anaerobic Hungate-type culture tubes (⌀16 x 125 mm, Glasgerätebau Ochs, Prod. No. 1020471) for transportation into the anaerobic chamber.

### Construction of a luminescent C. difficile reporter strain

A luminescent strain was constructed from *C. difficile* (Hall and O’Toole 1935) Lawson et al. 2016 strain 630 (DSM27543; NT5083), a virulent and multidrug-resistant strain (epidemic type X), which was isolated from a hospital patient with severe pseudomembranous colitis and had spread to several other patients on the same ward in Zurich, Switzerland ^47^. To obtain a luminescent *C. difficile* reporter strain expressing sLucOPT under the control of the constitutive *fdxA* promoter (CD630_01721), the sequence upstream of (and including) the *fdxA* transcription start site ^48^ (CP010905.2: 234479-234578, reverse strand) was PCR-amplified from *C. difficile* 630 genomic DNA using the S7 Fusion High-Fidelity Polymerase (Mobidiag, Prod. No. MD-S7-100), HF Buffer (Mobidiag, Prod. No. MD-B704), and oligos FFO-772/FFO-773 to append NheI- and SacI-restriction sites to the resulting PCR product. The PCR fragment and the sLucOPT-encoding vector pAP24 ^49^ were digested with FastDigest NheI (cat. No. FD0974) and FastDigest SacI (cat. No. FD1133), purified from agarose gel, and subsequently ligated using the T4 DNA ligase (Thermo-Fisher, cat. No. 15224017), resulting in pFF-189. The plasmid was transformed to *E. coli* TOP10 for propagation, transformed to the donor strain *E. coli* CA434 (HB101 carrying the IncPb conjugative plasmid R702), and finally delivered to *C. difficile* 630 (DSM 27543) by conjugation as described previously ^50^. The resulting plasmid-carrying strain, *C. difficile* [pFF-189], was designated FFS-515 (*i.e.*, LM0061 in Supplementary Table 1).

### In vitro *invasion assay for* C. difficile

To assess the ability of *C. difficile* to grow in drug- and/or pH-treated stool-derived and bioreactor communities, we used a luminescent-based assay, which we had already established before for *Gammaproteobacteria* ^29^. Here, we used the strain *C. difficile* LM0061. *C. difficile* LM0061 was grown anaerobically in mGAM containing 15 µg/mL thiamphenicol overnight and sub-cultured (1:100) in the same medium for another overnight culture before being used in the invasion assay.

### Stool-derived bacterial communities from healthy human donors

Drug master plates in DMSO were prepared as described before ^43^, with the difference that the lowest omeprazole concentration was omitted. Instead, row E only contained DMSO and served as a control. For clindamycin, the lowest two concentrations were omitted. A 96-well deep-well plate was prepared with 95 µL mGAM per well, and 5 µL of the drug master plate was transferred into it. These plates were stored frozen for a maximum of three weeks before being used. To test the effect of different pH on bacterial communities of human fecal samples on *C. difficile* growth, 96-well deep-well plates were prepared with 475 µL of mGAM at the respective pH inside the anaerobic chamber.

Glycerol stocks from human stool samples were prepared as previously described ^43^. Anaerobic overnight cultures from glycerol stocks were directly used for the assay. The drug-mGAM deep-well plates were pre-reduced in the anaerobic chamber for 24 h before being inoculated with 400 µL human fecal culture. Final drug concentration ranged from 2.5 µM to 160 µM for omeprazole and 10 µM to 100 µM for clindamycin with 1% DMSO and a starting human fecal OD_578_ of 0.01 per well. Wells containing communities from the same donor and 1% DMSO served as controls. The pH-mGAM plates were inoculated with 25 uL of the overnight culture from stool-derived communities to a final starting human fecal OD_578_ of 0.01 per well. Plates were grown for 24 h anaerobically at 37°C.

After the incubation, OD_578_ of every well was measured, and a fresh deep-well plate was prepared with 250 µL mGAM per well. Of the drug-treated- or pH-exposed human fecal samples, 50 µL were transferred into the fresh deep-well plate. This assay deep-well plate was used for pathogen challenge.

### Bioreactor communities

At every sampling time point, the OD_600_ of all bioreactors was measured, and a sample was transferred into a pre-reduced deep-well plate containing 250 µL mGAM (50 µL sample per well; 11 technical replicate wells per bioreactor; one plate per time point). This deep-well plate was used for pathogen challenge assay.

### Pathogen challenge (human fecal samples and bioreactor communities)

*C. difficile* LM0061 was diluted to an OD_578_ of 0.0025, and 200 µL were added to each well of the assay deep-well plate. The final volume was 500 µL (250 µL fresh mGAM, 50 µL drug-perturbed or pH-exposed fecal sample/bioreactor community, 200 µL *C. difficile*) and *C. difficile* starting OD_578_ was 0.001. The assay deep-well plate was sealed with an AeraSeal breathable membrane (Sigma-Aldrich, cat. No. A9224) and incubated at 37°C anaerobically for 5 h. After incubation, all wells were thoroughly mixed, and 100 µL per well were transferred to a white 96-well plate (Thermofisher 236105). This plate was brought out of the anaerobic chamber, and luminescence was measured with the Nano-Glo Luciferase Assay system kit from Promega (cat. No. N1110) in a Tecan Infinite 200 PRO microplate reader. The assay was done in three biological replicates.

For human fecal samples, the OD_578_ and luminescence were normalized to the values of the respective unperturbed controls (per donor and replicate), and the mean was calculated per drug concentration or pH, respectively.

For the bioreactor communities, the OD_600_ of every bioreactor was normalized to the median OD_600_ across all time points of the untreated control bioreactor. Luminescence data was analyzed per sampling time point. All values were normalized to the median of the untreated control bioreactor before taking the mean of all 11 technical replicates per condition and sampling time point.

### 16S rRNA gene amplicon sequencing

At every sampling point, 1 mL of the bioreactor cultures were harvested, and the pellets were frozen at −80°C for subsequent 16S rRNA gene amplicon sequencing. DNA extraction and sequencing were then conducted as described previously ^29^.

In brief, DNA was isolated with the DNeasy UltraClean 96 Microbial Kit (Qiagen 10196-4). Library preparation and sequencing were performed at the NGS Competence Center NCCT (Tübingen, Germany) with the 515F ^51^ and 806R ^52^ primers (covering a ~350-bp fragment of the 16S V4 region). Initial PCR products were purified, and indexing was performed in a second step PCR. After another bead purification, the libraries were checked for correct fragment length, quantified, and pooled equimolarly. The pool was sequenced on an Illumina MiSeq device with a v2 sequencing kit (input molarity 10 pM, 20% PhiX spike-in, 2×250 bp read lengths).

### Computational processing of 16S rRNA gene amplicon sequences

16S rRNA analysis was conducted using the *Dieciseis* R package from our lab, which uses the standard DADA2 workflow (https://benjjneb.github.io/dada2/bigdata.html). The *Dieciseis* pipeline is optimized for the analysis of our synthetic community Com21 and is derived from the workflow described in our previous work ^29^.

Briefly, quality profiles of the raw sequences were examined, trimmed, and paired-end reads filtered using the following parameters: trimLeft: 23, 24; truncLen: 225, 200; maxEE: 2, 2; truncQ: 11. The filtered forward and reverse reads were dereplicated separately, and amplicon sequence variants (ASVs) were inferred using default parameters. Subsequently, the reads were merged on a per-sample basis, and the merged reads were filtered to retain only those with a length between 244 and 245 bp before undergoing chimera removal.

Taxonomic assignment was carried out in two stages. First, the final set of ASVs was classified up to genus level using a curated DADA2-formatted database based on the genome taxonomy database (GTDB) release R06-RS202 ^53^ at https://scilifelab.figshare.com/articles/dataset/SBDI_Sativa_curated_16S_GTDB_database/14869077. Next, ASVs belonging to genera expected to be in Com21 were further classified at the species level using a modified version of the aforementioned database that contained only full-length 16S rRNA sequences of the 21 members of the synthetic community. The sequence of each ASV was aligned against this database using the R package DECIPHER v. 2.24.0 ^54^; we classified an ASV as a given species if it had sequence similarity >98% to the closest member in the database for 20/21 species. For *V. parvula* we had to change it to >95%. The abundance of each taxon of Com21 was obtained by aggregating reads at the species level.

## Data Availability

All 16S rRNA sequencing data generated in this study is available at the European Nucleotide Archive, accession ID PRJEB76870.

## Acknowledgments

The authors thank the NGS Competence Center Tübingen (NCCT, Germany). This work was supported by the Deutsche Forschungsgemeinschaft (DFG, German Research Foundation) under Grant EXC 2124 – 390838134 and Grant MA 8164/1-2, DZIF, and CEGIMIR.

## Author contributions

Conceptualization: B. M. and L. M.; Methodology: P. M., J. Sc. and J. Su.; Formal analysis: P. M. and J. Sc.; Investigation: P. M. and J. Sc.; Writing-Original Draft: P. M., J. Sc., B. M. and L. M.; Writing-Review & Editing: all; Supervision: F. F., B. M and L. M.; Funding Acquisition: F. F., B. M. and L. M.

## Declaration of interest

The authors declare no competing interests.

## Tables

**Supplementary Table 1.**
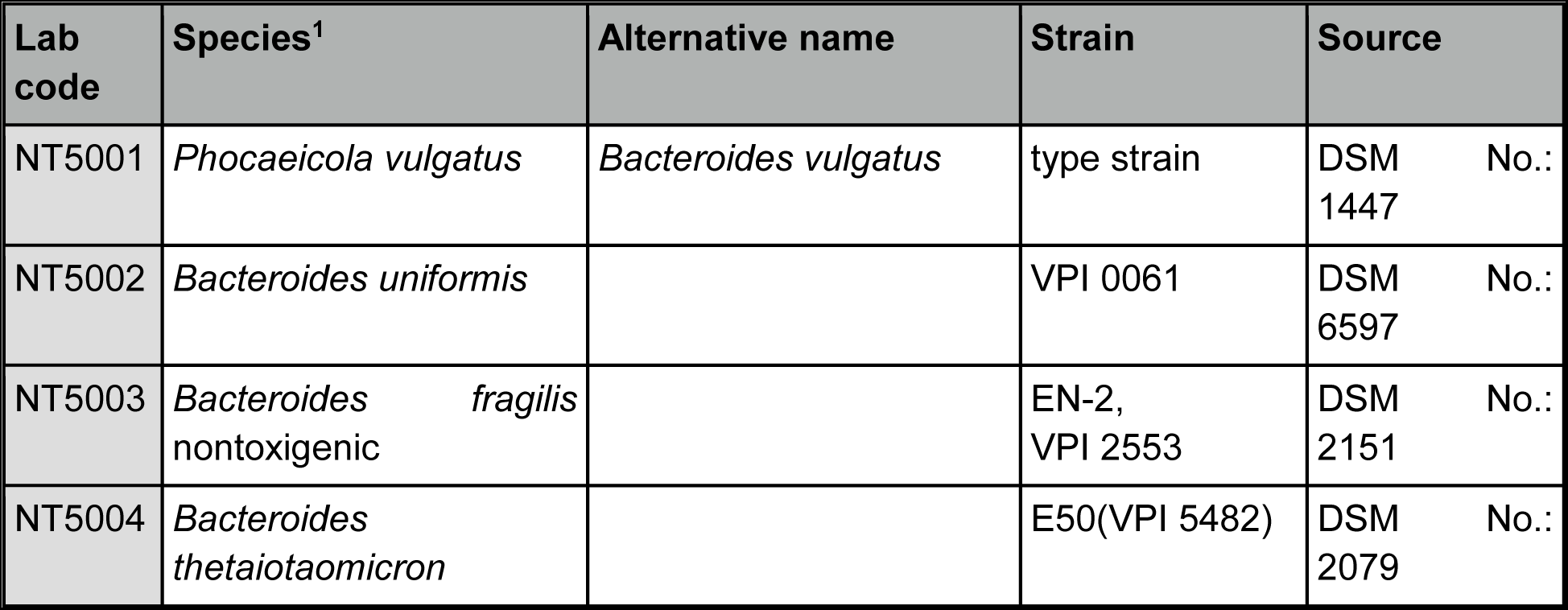
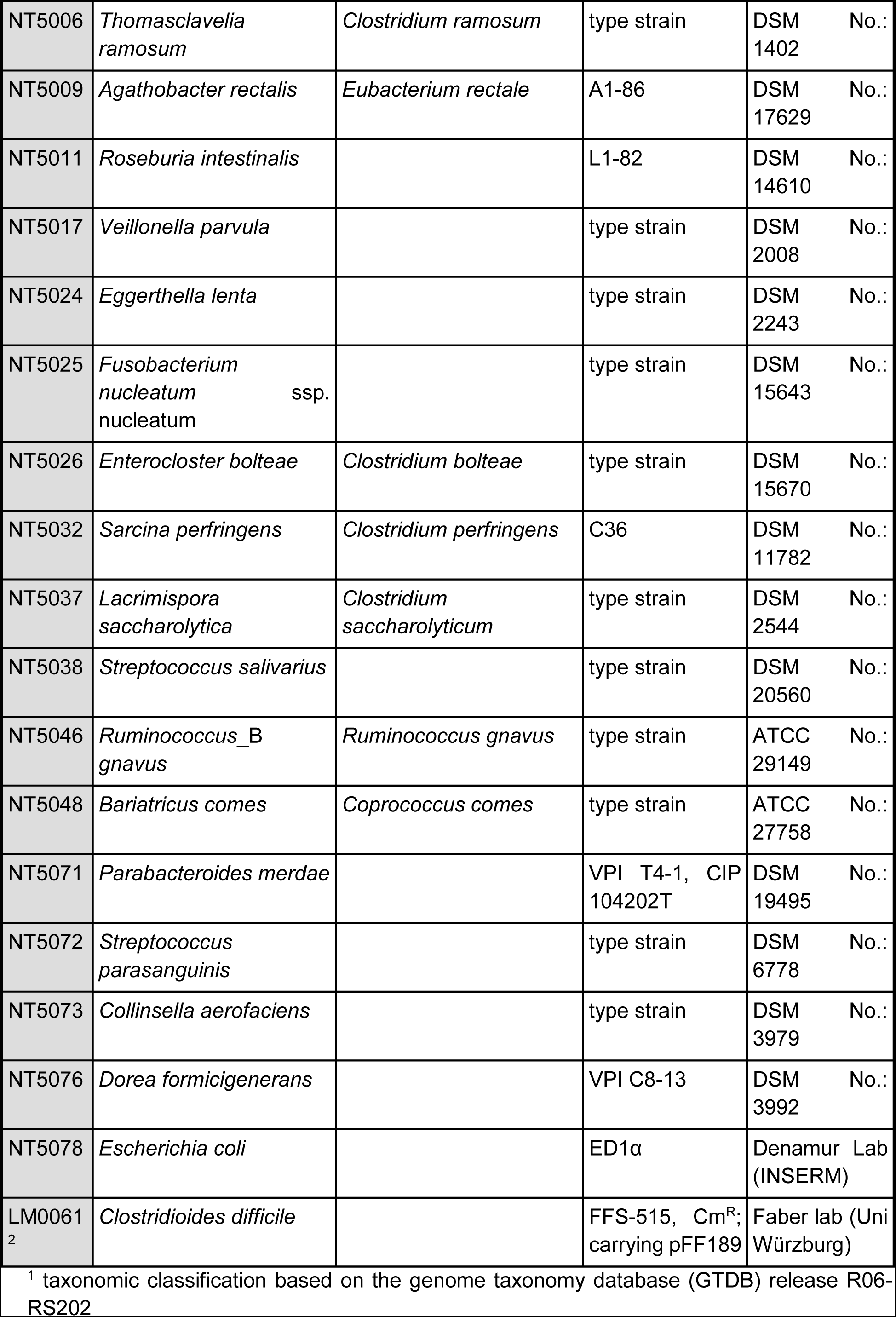
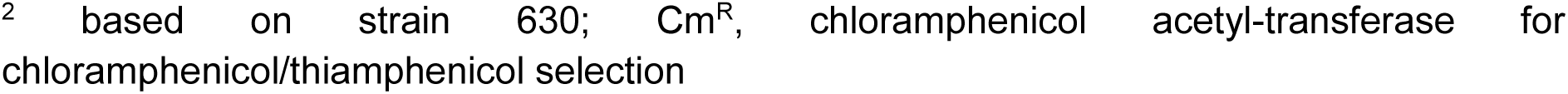
Strains used in this study.

**Supplementary Table 2.** Materials for the MBS.

| Item | Reference number | Supplier | Quantity |
| --- | --- | --- | --- |
| Heating thermostat (CC-104A) | 461-1056 | HUBER *via* VWR | 1 |
| Masterflex L/S® Multichannel Cartridge Pump Head with Reduced Pulsation for Microbore 2-Stop Tubing, 12-Channel, 8-Roller | HV-07519-25 | Cole-Parmer | 1 or 2 |
| Masterflex L/S® Small Cartridges for Multichannel Cartridge Pump Head with Reduced Pulsation for Microbore 2-stop Tubing | SI-07519-85 | Cole-Parmer | 12 |
| DURAN® double walled, wide mouth bottle GLS 80®, 500mL | 215-4156 | VWR | 6 |
| BOLA GLS 80 Vessel Closure \| PTFE \| 5 x GL 14 \| 1 x GL 25 | XZ019-182117 | Zinsstag | 6 |
| Multi-position magnetic stirrers, MIX series | 442-0752 | VWR | 1 |
| Masterflex L/S® Variable-Speed Digital Drive with Remote I/O, 1 to 100 rpm; 90 to 260 VAC | HV-07528-30 | Cole-Parmer | 1 or 2 |
| Masterflex C/L® Analog Variable-Speed Pump with Single-Channel Pump Head for Microbore Tubing Pump, 13 to 80 rpm; 90 to 260 VAC | 77122-32 | Cole-Parmer | 12 |
| MV5010 pH / Redox / ISE-Transducer with display in wall-mounting case | 90278080 | Xylem/Si-analytics | 6 |
| Item stand for MBS | Offer Nr.: AN00178860-1 | ITEM | 1 |
| Screws for MBS system | 6834886 | Hornbach | 1 pck |
| Power strip | 3882931 | Hornbach | 2 |
| Tygon tubing A-60-G, I/P 73, 50 ft. | MFLX06404-73 | VWR | 2 |
| Flexible Cable H05VV-F 4 x 1 mm<sup>2</sup>, black, sold by meter | 1499067 | Conrad | 10m |
| Ferrule 1 mm<sup>2</sup> Partially Insulated | 617836 - 62 | Conrad | 1 pck |
| Octagon 8 port - manifold | 343938 | Huber | 1 |
| Masterflex® Ismatec® Pump Tubing, 2-Stop, Viton®, 2.06 mm ID, 15" L; 12/PK | MFLX96428-42 | VWR | 1 |
| Laboratory Screw Joints, GL14 cap, 3-parts, including PTFE/ETFE fittings (6mm) | D590-06 | Bola | 30 |
| Laboratory Screw Joints, GL25 cap, 3-parts, including PTFE/ETFE fittings (12mm) | D590-34 | Bola | 6 |
| Hose connectors ROTILABO® Y-shape with conical ends, hose inner Ø 9-11mm | TT53.1 | Carl Roth | 1 pck |
| Hose connectors ROTILABO® T-shape, Hose inner Ø 10-11 mm | E767.1 | Carl Roth | 1 pck |
| Hose connectors ROTILABO® angle shape, Hose inner Ø 10-11 mm | E788.1 | Carl Roth | 1 pck |
| Rapid couplings, male connections, hose nipples with hose nozzles, Ø9.5mm | 8771-1060 | Bürkle | 8 |
| ATEX II 1/2G pH-single-rod measuring cell | SL 81-225 pH<br>VP | Xylem/Si-analytics | 6 |
| Variopin cable | 85442000 | Xylem/Si-analytics | 6 |
| Disposable needles Sterican® long bevel facet, 30 mm, 0.60 mm, blue | X129.1 | Carl Roth | 6 per experiment |
| Button Once Canulla, 45 mm length | KK45R21S | CLS<br>Medizintechnik und Vertrieb | 6 per experiment |
| Gas trap | 114100 | Ochs | 6 |
| Pump Tubing, PharMed® BPT, 1,14 mm ID; 100 ft | MFLX95809-30 | VWR | 1 pck |
| Masterflex L/S Norprene Food-Grade Tubing, L/S 16, 50ft. | MFLX06402-16 | VWR | 2 pck |
| Magnetic bars ROTILABO® Economy, Ø: 8 mm, 25 mm | XA18.1 | Carl Roth | 6 |
| Set of customized stainless-steel tubing, Ø 6mm, tube (240 mm) with sparger, sample tubing (200 mm), off-gas tubing (100 mm) | Offer Nr.: BZV-2022111600511 | bbi-biotech | 6 of each |
| 5 L Tedlar® Gas sampling bags, 2-in-1 PP valve Thermogreen® LB-2 Septa | 24655 | VWR | 1 |
| J.T.Baker®, Syringe Filters, pore size 0,22 µm | SF02-60 | VWR | 100 |
| Luer/Lock fittings in different sizes | CT59.1-64.1 | Carl Roth | 150 |
| 5-L duran bottle | 215-0057 | VWR | 2 |
| 1-L duran bottle | 215-1595 | VWR | 2 |

## Figure Legends

**Extended Data Figure 1.**
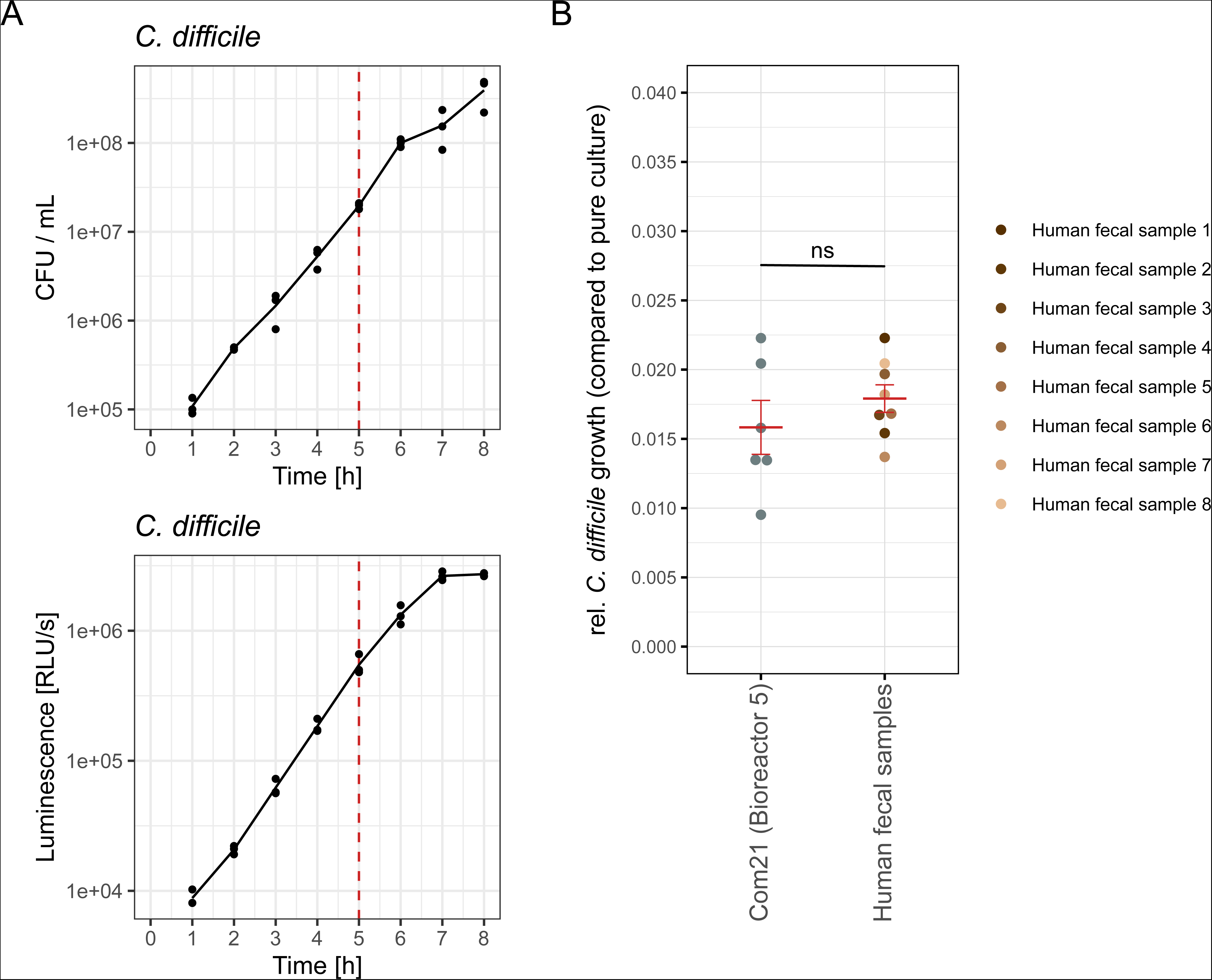
**A)** Growth curves for *C. difficile* LM0061 based on plating on mGAM agar (top) or luminescence (bottom). The lines indicate the mean of three biological replicates. Red vertical lines mark the endpoint of the *C. difficile* invasion assay, which falls within the linear range of the curves, allowing luminescence to be used as a proxy for *C. difficile* levels. **B)** Relative growth of *C. difficile* during co-culture with untreated Com21 from bioreactor 5 or human fecal samples compared to pure culture. Pathogen levels were quantified via luminescence after 5 h. For Com21 values are shown from each sampling point of the multiple-bioreactor system (six in total) from bioreactor 5 (pH 7, untreated). For human fecal samples the mean of three biological replicates is shown per fecal sample. Red points and bars represent mean (M) ± standard error of the mean (SEM). No significant (ns) difference in relative *C. difficile* growth between Com21 from bioreactor 5 (M = 0.0158, SEM = 0.0019) and human fecal samples (M = 0.0179, SEM = 0.001). Two-sided t-test: t(12) = −1.026; p = 0.3251; d = 0.5541.

**Extended Data Figure 2.**
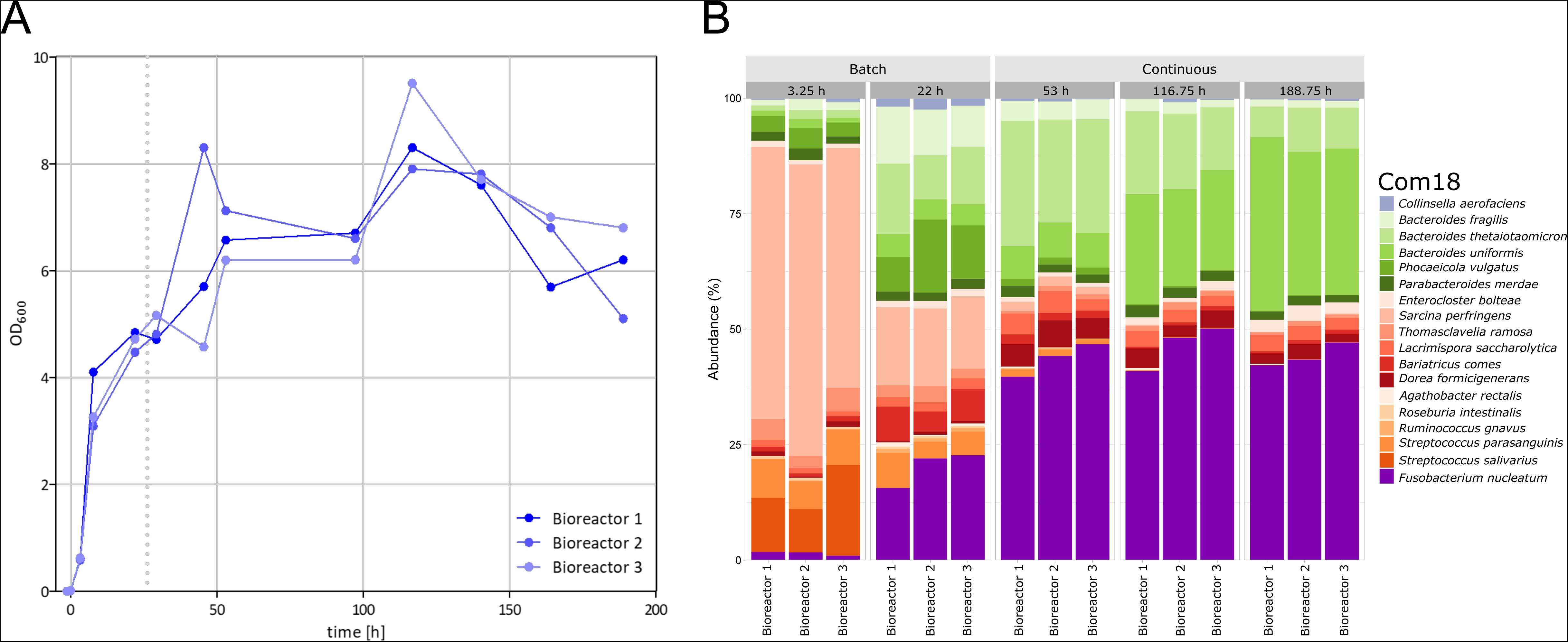
Continuous growth of Com18 in the MBS. **A)** OD of the bioreactors throughout the entire operation period. The community was grown in batch mode for one day before switching to continuous mode (indicated by the gray dashed line). **B)** Relative abundance of each strain in the Com18 at the indicated time points. Triplicates are presented in one panel. The first two sampling time points were during batch mode, while the subsequent sampling time points were during continuous mode.

## Notes

### Competing Interest Statement

The authors have declared no competing interest.

